# Decoding the Moving Mind: Multi-Subject fMRI-to-Video Retrieval with MLLM Semantic Grounding

**DOI:** 10.1101/2025.04.07.647335

**Authors:** Xuanliu Zhu, Yiqiao Chai, Runnan Li, Mingying Lan, Li Gao

## Abstract

Decoding dynamic visual information from brain activity remains challenging due to inter-subject neural heterogeneity, limited per-subject data availability, and the substantial temporal resolution gap between fMRI signals (0.5 Hz) and video dynamics (30 Hz). Current approaches face persistent challenges in addressing these temporal mismatches, demonstrate limited capacity to integrate subject-specific neural patterns with shared representational frameworks, and lack adequate semantic granularity for aligning neural responses with visual content. To bridge these gaps, we propose a framework addressing these limitations through three innovations: (1) a Dynamic Temporal Alignment module that resolves temporal mismatches via exponentially weighted multi-frame fusion with adaptive decay coefficients; (2) a Brain Mixture-of-Experts architecture that combines subject-specific extractors with shared expert layers through parameter-efficient tri-modal contrastive learning; and (3) a Multi perspective Semantic Hyper-Anchoring module that resolves cross-subject attention bias via multi-dimensional semantic decomposition, leveraging multimodal LLMs for fine-grained video semantic extraction—enabling the model to match individual attention patterns as different subjects naturally focus on distinct aspects of the same visual stimulus. This module boosts Top-10/Top-100 retrieval by 17.7%/6.6%. Experiments on two video-fMRI datasets demonstrate **state-of-the-art** performance, with **39%/30%** improvements in Top-10/Top-100 accuracy over single-subject baselines and 27% gains against multi-subject models. The framework exhibits remarkable few-shot adaptability, retaining **97%** performance when using only **10%** training data for new subjects. Visualization analysis confirms this generalization capability stems from effective disentanglement of subject-specific and shared neural representations.

## 1 Introduction

In recent years, functional magnetic resonance imaging (fMRI)-based visual decoding has made significant progress, with researchers gradually achieving decoding from low-level visual features to high-level semantics. Neural decoding approaches include classification [1, 2, 3], retrieval [4, 5], and reconstruction [6, 7], with this study focusing on retrieval tasks. Previous studies have achieved notable success in retrieving static stimulus images [4, 5]. However, since most visual stimuli encountered in daily life are continuous and dynamic, video retrieval from brain signals holds greater practical value.

Nevertheless, dynamic video retrieval for natural visual experiences faces three core challenges: First, there exists a temporal resolution gap of tens of times between the slow hemodynamic response (∼ 0.5 Hz) of blood-oxygen-level-dependent (BOLD) signals [8, 9] and video temporal dynamics (30 Hz). In prior work, Lu et al. [10] employed multilayer perceptrons to extract fMRI features but neglected the temporal lag effect of BOLD signals. Chen et al. [11] considered information across multiple fMRI frames, yet the low signal-to-noise ratio of fMRI may lead to overfitting when using Transformer [12] to extract temporal features from adjacent frames.

Second, due to inter-subject variability in brain structure and function [13], most existing decoding models are subject-specific [14, 15, 16, 17], limiting their generalizability. The scarcity of fMRI data for new subjects often leads to overfitting, as single-subject datasets are typically small. Multi-subject decoding methods address these issues by pooling data across individuals, enhancing generalization, reducing training costs, and improving computational efficiency. Thus, developing such methods is critically important.

Moreover, conventional semantic feature extraction methods predominantly rely on image caption models [18, 19, 20], which struggle to capture fine-grained semantics of dynamic visual content and are prone to semantic granularity loss [11] in complex dynamic scenes.

In summary, existing cross-subject video decoding models face the following challenges: (1) **The temporal resolution gap** between slow fMRI hemodynamic responses (0.5 Hz) and video dynamics (30 Hz); (2) **Functional regional distribution and response characteristics vary across subjects**, making it difficult for existing methods to simultaneously extract subject-specific and shared information, resulting in low parameter efficiency; (3) Traditional semantic alignment methods rely on shallow text embeddings, **failing to capture nuanced visual semantics in dynamic scenes**.

To address these challenges, we propose **CrossMind-VL**, a novel method for cross-subject brain decoding using a single model: (1) **Dynamic Temporal Alignment**: We propose an exponentially weighted multi-frame fusion module to align fMRI’s hemodynamic delays (0.5 Hz) with video dynamics (30 Hz).

Adaptive decay coefficients aggregate delayed fMRI signals, preserving video temporal continuity. (2) **BrainMoE Architecture**: To bridge functional heterogeneity across subjects, we propose a Brain-based Mixture-of-Experts (BrainMoE) architecture that separates subject-specific feature projection from shared semantic alignment. Subject-specific layers extract unique information for each subject, while shared layers capture common patterns across subjects, with final alignment to a unified feature space via graph-text-brain tri-modal contrastive learning. (3) **Multi-perspective Semantic Hyper-Anchoring**: We develop a fine-grained video semantic extraction framework using multimodal large language models (MLLMs), which addresses cross-subject attention bias in neural decoding through multi-dimensional semantic decomposition. Ablation studies demonstrate that this module significantly improves Top-10/Top-100 retrieval accuracy by 17.7% and 6.6%, effectively mitigating the “selective attention paradox” caused by cognitive focus variations. This enhancement validates the critical role of fine-grained semantic representation in establishing robust fMRI-video cross-modal correspondences.

Our key contributions are summarized as follows:

- **First introduction of MLLM to fMRI retrieval**: We leverage Qwen2-VL [21] to achieve human-level semantic understanding of dynamic visual content, resolving the “semantic granularity bottleneck” in prior work.
- **Multi-frame fusion module**: We propose an exponentially weighted multi-frame fMRI fusion mechanism specifically designed for hemodynamic lag characteristics, significantly enhancing fMRI representation capability.
- **BrainMoE architecture**: The first feature extractor combining subject-specific modules with dynamic gated MoE layers, requiring only 24% parameter fine-tuning for unseen subjects while improving cross-subject generalization by 54%.
- **State-of-the-art performance**: Achieves 39% and 30% improvements in Top-10 and Top-100 accuracy respectively over previous single-subject SOTA models on two public video-fMRI datasets.

## 2 Related Work

Neural decoding [22, 23, 24, 25] constitutes the computational process of extracting perceptual information from neuroimaging signals, with methodological frameworks stratified by task complexity into classification, retrieval, and reconstruction. This work focuses primarily on visual retrieval paradigms.

### 2.1 Static Image Decoding

Systematic methodologies have been established for fMRI-based static image decoding. Haxby et al. [26] pioneered multivoxel pattern analysis (MVPA) to achieve eight-category visual stimulus classification, empirically validating the multivariate discriminability of neural representations. For primary visual feature extraction, Kamitani et al. [27] demonstrated the efficacy of linear support vector machines (SVMs) in decoding low-order features (e.g., orientation selectivity), though constrained by linear assumptions in capturing high-level semantic representations. Deep learning advancements have catalyzed semantic space mapping frameworks: Stansbury et al. [28] developed latent Dirichlet allocation (LDA) models for scene co-occurrence statistics, while Li et al. [24] proposed multi-label deep networks to resolve complex semantic concurrency. Breakthroughs in image retrieval emerged through neural encoding models [29] and CNN-based feature decoders [2], establishing cross-modal mappings between brain signals and image feature spaces.

### 2.2 Dynamic Video Decoding

Despite progress in static visual decoding, dynamic visual information processing confronts fundamental challenges: naturalistic visual stimuli inherently exhibit spatiotemporal coupling, while the temporal resolution disparity between fMRI (sampling interval ≈ 2 s) and video streams (typically 30 fps) spans three orders of magnitude. Nishimoto et al. [17] pioneered motion energy modeling combined with Bayesian regression for video retrieval. Subsequent approaches by Han [30] and Wen et al. [31] employed dimensionality reduction to single-frame AlexNet features with linear regression. Lu et al. [10] implemented multilayer perceptrons to extract fMRI features aligned with Contrastive Language–Image Pre-training (CLIP) [32] embeddings via contrastive learning, albeit neglecting temporal latency effects in BOLD signals. Notably, Chen et al. [33] introduced temporal feature extraction through transformer architectures across adjacent fMRI frames. Building upon this foundation, our approach incorporates multi-frame fMRI signals but diverges by implementing exponentially decaying weights for signal integration—a design choice mitigating overfitting risks associated with low signal-to-noise fMRI data.

### 2.3 Cross-Subject Decoding

Current decoding models predominantly operate in single-subject regimes, constrained by data scarcity and cross-subject generalization barriers. The core challenge stems from inter-individual neural heterogeneity: while anatomical variations can be mitigated through image registration, functional topographical variability—encompassing spatial distributions and response properties of functional regions—remains a fundamental obstacle [34].

Existing alignment methods exhibit critical limitations: Hyperalignment projects neural responses into shared semantic spaces [34] but falters with stimulus content mismatches; Shared Response Models (SRMs) decompose subject-specific bases and shared responses [35] yet require strict temporal alignment; Graph decoding models (GDMs) construct cross-subject similarity graphs for feature alignment [36] but suffer from computational complexity and graph construction sensitivity.

To address these limitations, we propose a novel Mixture-of-Experts (MoE) [37] framework for cross-subject decoding, achieving dual breakthroughs: 1) A hierarchical decoding architecture employing gating networks to dynamically integrate shared experts and private experts; 2) A sparse activation mechanism maintaining parameter efficiency independent of cohort size while enhancing computational scalability. This formulation explicitly addresses the tripartite challenge of neural heterogeneity, data scarcity, and model generalizability.

## 3 Method

### 3.1 Problem Statement

Given a neural dataset comprising neural activities of *N* subjects under visual stimuli, let **X**^(*n*)^ denote the voxel space for the *n*-th subject. Here, 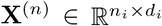 represents the neural responses of the *n*-th subject, where *n*_*i*_ is the number of image stimuli and *d*_*i*_ is the number of voxels. Let **I** and **Y** denote the pixel space and the label space of the image stimuli, respectively.

Our objective is to align the fMRI representations of the *N* subjects into a unified representation space and perform zero-shot retrieval tasks. To achieve this, we first leverage a multimodal large language model (MLLM) to obtain semantically rich labels for the visual stimuli. Subsequently, we utilize the CLIP visual and text branches to extract visual and semantic features, respectively. We then propose a BrainMOE architecture to separately extract subject-specific features and subject-shared features. Finally, we align this feature space with the CLIP representation space through tri-modal contrastive learning.

### 3.2 Feature Extraction

#### 3.2.1 Video Feature Extraction

To extract visual features from video stimuli, we follow a frame-based approach. For each video segment **V**_*i*_ of duration 2 seconds, we uniformly sample *T* = 8 frames, denoted as 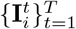, where 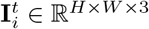 represents the *t*-th frame. Each frame is processed through the pre-trained CLIP image encoder to obtain its visual feature representation. The final visual feature 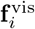 for the video segment is computed as the average of the frame-level features:

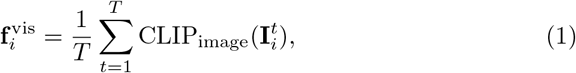

where CLIP_image_(·) denotes the pre-trained CLIP image encoder. This approach ensures that the temporal information in the video is captured while maintaining computational efficiency.

#### 3.2.2 Fine-grained Semantic Label Acquisition

To generate fine-grained semantic labels for video stimuli, we employ the Qwen2-VL (7B) model, a state-of-the-art multimodal large language model capable of understanding video content. For each video segment **V**_*i*_, we input the prompt: “Describe the content of this video from five different perspectives.” The model generates five textual descriptions 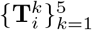, where 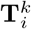 represents the *k*-th description. These descriptions capture diverse aspects of the video content, such as objects, actions, and context.

Each textual description 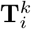 is encoded using the pre-trained CLIP text encoder to obtain its semantic embedding. The final semantic feature 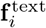 is computed as the average of these embeddings:

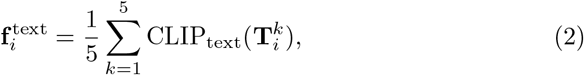

where CLIP_text_(·) denotes the pre-trained CLIP text encoder. This method ensures that the semantic representation of the video is both rich and comprehensive.

### 3.3 Enhancement of fMRI Temporal Dynamic Information

Given a training set 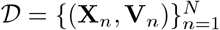, where **X**_*n*_ ∈ ℝ^*D*^ represents the fMRI data of the *n*-th subject (preprocessed as a 1D vector of voxels) and **V**_*n*_ denotes the corresponding video stimuli, we address the significant temporal resolution gap between fMRI data (sampled every 2 seconds) and video frames (30 frames per second). To bridge this gap and enrich the temporal dynamics of the fMRI data, we follow the approach of Chen et al. [11] and propose a temporal interpolation method. Unlike their method, which employs cross-frame attention, we directly use exponential weighting to reduce computational cost.

The key idea is to extend each fMRI frame **X**_*n*_ to a sequence of *W* +1 frames, where *W* is the size of the time window. The interpolated sequence is defined as:

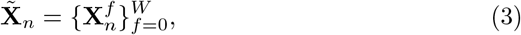

where 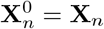 is the original frame. The interpolated frames 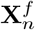 for *f >* 0 are computed using an exponential interpolation function:

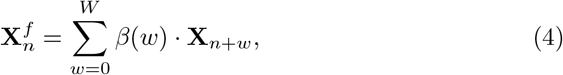

where *β*(*w*) represents the interpolation weights, defined as:

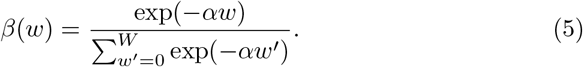

Here, *α* controls the rate of exponential decay (set to *α* = 0.1 in practice), reflecting the diminishing influence of distant frames. The normalization condition 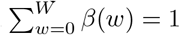 ensures that the overall signal magnitude is preserved. The parameters *W* and *α*, which determine the size of the time window and the rate of exponential decay, respectively, are experimentally validated in Section 4.7.

This interpolation method effectively models the temporal dynamics of fMRI data, providing a richer representation that aligns more closely with the continuous flow of visual information in the video stimuli.

### 3.4 Training Pipeline of the BrainMOE

#### 3.4.1 BrainMOE Module

The BrainMOE module consists of two main components: the **Subject-Specific Layer** and the **Subject-Shared Layer**. Below, we provide a formal description of each component.

##### Subject-Specific Layer

For each subject *n*, a dedicated linear layer is assigned to extract subject-specific features. Given the fMRI data 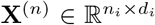 of the *n*-th subject, the subject-specific feature 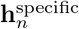 is computed as:

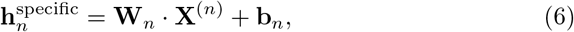

where 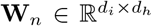 and 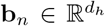 are the learnable weight matrix and bias vector for the *n*-th subject, respectively. **During training, only the parameters of the corresponding linear layer are updated using the data from the** *n***-th subject**.

##### Subject-Shared Layer

The Subject-Shared Layer is designed to extract shared features across all subjects. It consists of two residual blocks, each preceded by a Layer Normalization (Layer Norm) operation. The first residual block contains a Multi-Layer Perceptron (MLP), while the second residual block incorporates a Mixture of Experts (MOE) module with an MLP backbone. Formally, the shared feature **h**^shared^ is computed as follows:

First, the input **h**_0_ (e.g., the concatenation of subject-specific features) is processed through the first residual block:

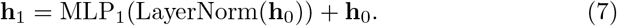

Next, the output of the first residual block **h**_1_ is processed through the second residual block:

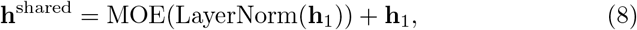

where the MOE module is defined as:

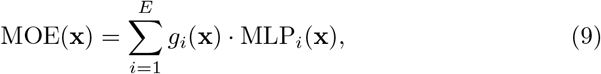

with *E* experts, *g*_*i*_(**x**) as the gating function for the *i*-th expert, and MLP_*i*_(**x**) as the MLP backbone for the *i*-th expert.

The parameters of the Subject-Shared Layer are updated using data from all subjects, ensuring the extraction of common features across the population.

#### 3.4.2 Tri-Modal Contrastive Learning with BrainMOE

To align the fMRI representations with the visual and semantic features extracted from CLIP, we employ a tri-modal contrastive learning framework. The loss function consists of two components: the projection loss *L*_*Projection*_ and the semantic loss *L*_*Semantic*_.

The projection loss *L*_*Projection*_ ensures the alignment between fMRI embeddings **f**_*i*_ and text embeddings **t**_*i*_:

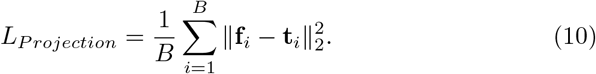

The semantic loss *L*_*Semantic*_ is the tri-modal contrastive loss, computed as:

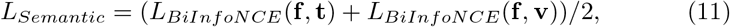

where *L*_*BiInfoNCE*_ is the bidirectional InfoNCE loss defined as:

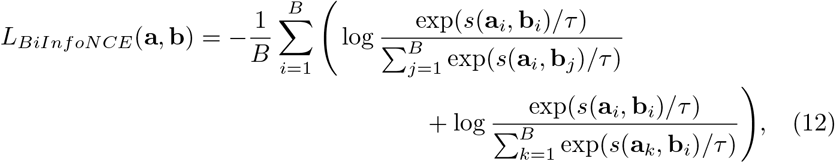

with *s*(·,·) denoting cosine similarity, *τ* as the temperature parameter, and *B* as the batch size.

The combined loss function is defined as:

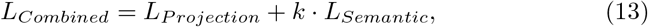

where *k* is a hyperparameter balancing the two losses.

This training pipeline leverages the BrainMOE module to extract both subject-specific and shared features, while the tri-modal contrastive learning aligns the fMRI representations with the visual and semantic spaces of CLIP.

## 4 Experiments

### 4.1 Datasets

In this study, we employ two publicly available video-fMRI datasets, each containing paired stimulus videos and corresponding fMRI responses from multiple healthy subjects (Table 1).

**Table 1:** Characteristics of the video-fMRI datasets used in our experiments.

| Dataset | Adopted subjects | TR | Train samples | Test samples |
| --- | --- | --- | --- | --- |
| CC2017 ([31]) | 3 | 2s | 4320 | 1200 |
| HCP ([38]) | 3 | 1s | 2736 | 304 |

#### CC2017

The CC2017 dataset [31] includes fMRI recordings from three subjects viewing movie clips, with 18 training and 5 testing movies (8 minutes each). Data were acquired using a 3T MRI system and preprocessed with standard steps including motion correction and MNI space registration. We identified visually activated voxels by computing time series correlations across training trials, applying Fisher’s z-transform, and selecting the top 4500 voxels via paired t-test. BOLD signals were adjusted for a 4-second hemodynamic delay [17, 39].

#### HCP

From the Human Connectome Project [38], we analyzed data from three representative subjects (100610, 102816, 104416) acquired using a 7T scanner (1.6 mm resolution). Preprocessing included motion/distortion correction and ICA-based artifact removal. Using the Glasser parcellation [40], we extracted 5820 voxels from visual ROIs (V1-V4), with identical 4-second hemodynamic delay correction as CC2017. The video segments were split into 90% training and 10% testing sets, following established practices [41].

### 4.2 Baseline Methods

To evaluate the effectiveness of the proposed approach, we conducted comparisons with both state-of-the-art single-subject decoding techniques and multisubject decoding methods.

#### Single-subject methods

The video decoding models proposed by Wen [31], Kupershmidt [42], Mind-video [11], and Mind-animator [10] are all trained on single-subject data. Specifically, Lu et al. [10] have evaluated theses retrieval performance on the CC2017 dataset.

#### Multi-subject methods

To the best of our knowledge, this is the first work to explore multi-subject video decoding. To evaluate the model’s performance in this context, we aggregated data from multiple subjects and trained both MLP and Transformer models on the combined dataset. We refer to these baseline methods as MS-MLP and MS-Transformer.

### 4.3 Evaluation Metrics

This study focuses on the retrieval task, which involves indexing the corresponding stimulus video from a predefined datapool based on the elicited fMRI signal. Compared to classification tasks, this task demands finer feature granularity and better reflects model performance. We employ top-10 and top-100 accuracy as evaluation metrics. To evaluate the model’s generalization capability, retrieval is performed not only on the original test set (“Small”) but also on an extended stimulus set (“Large”). For instance, in the CC2017 dataset, the “Small” version involves retrieval within the dataset’s test videos, while the “Large” version expands the datapool by incorporating another 3040 videos.

### 4.4 Hyperparameter settings

We trained **BrainMoE** on all datasets using consistent hyperparameters: a 3-frame fMRI aggregation window with exponential decay (*α* = 0.1), batch size of 256, and 25 training epochs optimized by Adam (learning rate=5 × 10^−5^). The architecture incorporated both subject-specific 768-dimensional linear projections and cross-subject representations through a 4-expert Mixture of Experts (MoE) system, where each expert utilized 24-dimensional hidden layers in its MLP. The complete training process converged within approximately one hour on a single NVIDIA 4090Ti GPU.

### 4.5 Comparative experiments results

Our experiments were conducted on both CC2017 and HCP datasets, with quantitative results presented in Tables 2 and 3, and qualitative demonstrations shown in Figure 2. Overall, our BrainMOE achieved state-of-the-art (SOTA) performance on both datasets. Under the ‘Small’ setting of the CC2017 dataset, the average inter-subject top-10 and top-100 accuracy improved by **39%** and **30%**, respectively, compared to the best single-subject model, Mind-Animator, and by **27%** and **6%** compared to MS-Transformer. Under the ‘Large’ setting, the average inter-subject top-10 and top-100 accuracy improved by **35%** and **16%** over Mind-Animator and by **12%** and **9%** over MS-Transformer. On the HCP dataset under the ‘Small’ setting, the average inter-subject top-10 and top-100 accuracy improved by **14%** and **7%** over MS-Transformer, while under the ‘Large’ setting, the improvements were **7%** and **13%**, respectively. Figure 2 demonstrates that BrainMoE achieves precise retrieval performance, accurately identifying the viewed video segment from semantically similar candidates based solely on brain activity patterns.

**Table 2:** Retrieval accuracy (%) on CC2017 dataset. For the “small test set”, the chance-level accuracies for top-10 and top-100 accuracy are 0.83% and 8.3%, respectively. For the “large test set”, the chance-level accuracies for top-10 and top-100 accuracy are 0.24% and 2.4%, respectively.

| Dataset | Number of models | Comparison methods |  | Subject1 |  | Subject2 |  | Subject3 |  | Average |  |
| --- | --- | --- | --- | --- | --- | --- | --- | --- | --- | --- | --- |
|  |  | Model | Test set | top-10 | top-100 | top-10 | top-100 | top-10 | top-100 | top-10 | top-100 |
| CC2017 | 3 | Wen [31] [Cerebral Cortex 2018] | Small | 2.17 | 19.50 | 3.33 | 19.17 | — | — | 2.75 | 19.33 |
|  |  | Kupersmidt [42] |  | 1.09 | 8.57 | 0.92 | 8.24 | 0.84 | 8.24 | 0.95 | 8.35 |
|  |  | Mind-video [11] [NeurIPS 2023 Oral] |  | 3.22 | 19.08 | 2.75 | 16.83 | 3.58 | 22.08 | 3.18 | 19.33 |
|  |  | Mind-animator [10] [ICLR 2025] |  | 3.08 | 22.58 | 4.75 | 26.90 | 4.50 | 24.67 | 4.11 | 24.72 |
|  | 1 | MS-MLP |  | 3.33 | 20.83 | 3.50 | 23.25 | 3.66 | 24.41 | 3.50 | 22.83 |
|  |  | MS-Transformer |  | 3.75 | 26.58 | 4.66 | 32.75 | 5.16 | 32.00 | 4.52 | 30.44 |
|  |  | Ours |  | **5.00** | **27.92** | **6.17** | **34.83** | **6.00** | **33.92** | **5.72** | **32.22** |
|  | 3 | Wen [31] [Cerebral Cortex 2018] | Large | 1.41 | 11.58 | 2.08 | 9.58 | — | — | 1.75 | 10.58 |
|  |  | Kupersmidt [42] |  | 0.17 | 2.94 | 0.17 | 2.77 | 0.25 | 2.18 | 0.19 | 2.63 |
|  |  | Mind-video [10] [NeurIPS 2023 Oral] |  | 1.75 | 7.17 | 0.83 | 5.17 | 1.25 | 9.00 | 1.28 | 7.11 |
|  |  | Mind-animator [10] [ICLR 2025] |  | 2.17 | 12.50 | 2.25 | 17.00 | 2.75 | 16.42 | 2.39 | 15.31 |
|  | 1 | MS-MLP |  | 2.08 | 10.00 | 1.66 | 11.16 | 2.66 | 14.91 | 2.13 | 12.02 |
|  |  | MS-Transformer |  | 2.08 | 13.58 | 3.08 | 17.33 | **3.58** | 17.66 | 2.91 | 16.19 |
|  |  | Ours |  | **2.67** | **14.33** | **3.58** | **19.42** | 3.50 | **19.33** | **3.25** | **17.69** |

**Table 3:** Retrieval accuracy (%) on HCP dataset. For the “small test set”, the chance-level accuracies for top-10 and top-100 accuracy are 3.29% and 32.9%, respectively. For the “large test set”, the chance-level accuracies for top-10 and top-100 accuracy are 0.66% and 6.6%, respectively.

| Dataset | Number of models | Comparison methods |  | Subject1 |  | Subject2 |  | Subject3 |  | Average |  |
| --- | --- | --- | --- | --- | --- | --- | --- | --- | --- | --- | --- |
|  |  | Model | Test set | top-10 | top-100 | top-10 | top-100 | top-10 | top-100 | top-10 | top-100 |
| HCP | 1 | MS-MLP | Small | 10.53 | 58.88 | 11.18 | 54.61 | 11.84 | 60.86 | 11.18 | 58.11 |
|  |  | MS-Transformer |  | 11.84 | 76.97 | 11.18 | 74.67 | 11.51 | 77.30 | 11.51 | 76.32 |
|  |  | Ours |  | **13.49** | **82.24** | **13.16** | **79.61** | **12.83** | **82.24** | **13.16** | **81.36** |
|  |  | MS-MLP | Large | 3.95 | 21.38 | 4.28 | 18.42 | 3.62 | 24.01 | 3.95 | 21.27 |
|  |  | MS-Transformer |  | 6.25 | 39.14 | 6.58 | 38.49 | **6.58** | 40.46 | 6.47 | 39.36 |
|  |  | Ours |  | **7.24** | **44.41** | **6.91** | **42.76** | **6.58** | **45.72** | **6.91** | **44.30** |

**Figure 1:**
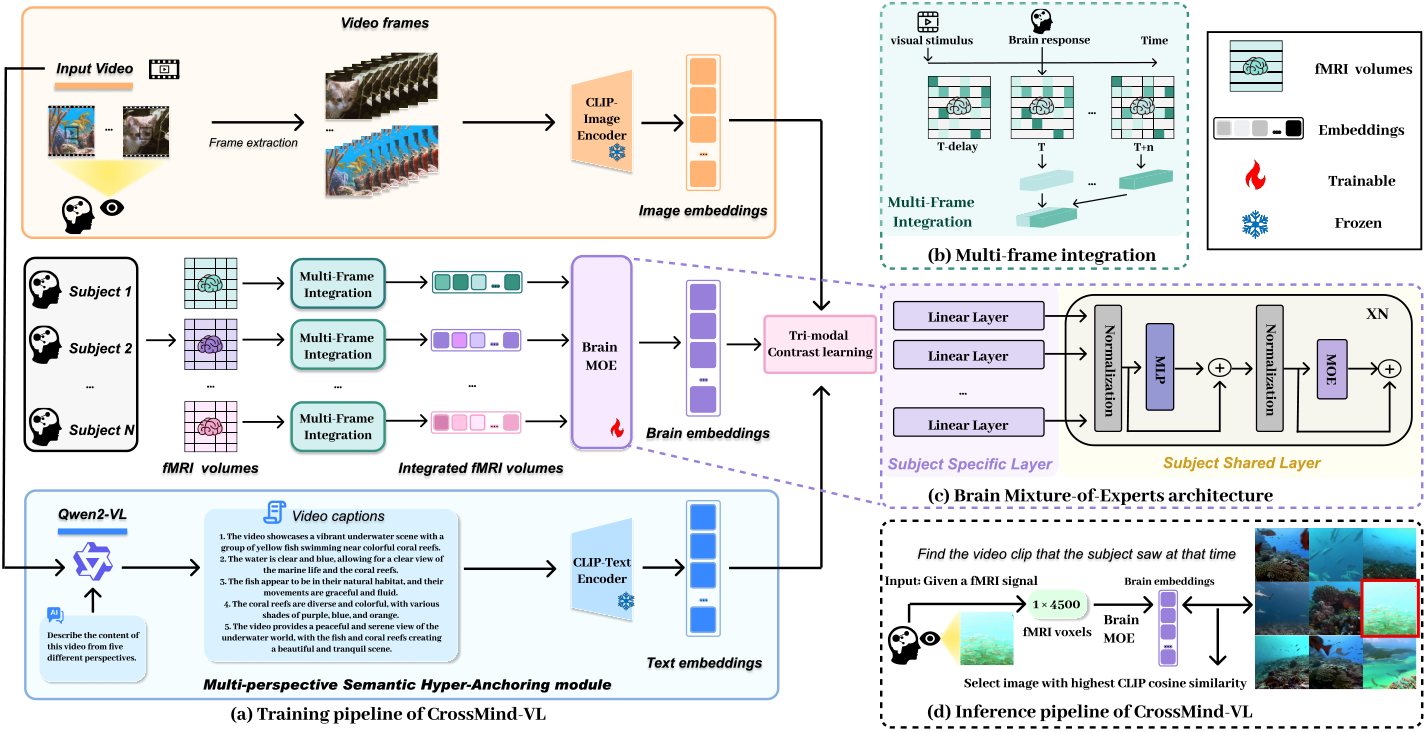
The framework of the proposed CrossMind-VL. (a) Training Pipeline of CrossMind-VL. Initially, Qwen2-VL is employed to annotate the stimulus video content in detail. Subsequently, CLIP is utilized to extract image and text features from the videos. Finally, contrastive learning is applied to align the features of the three modalities—image, text, and brain signals—into a unified representational space. (b) Multi-frame integration module. We aggregate fMRI signals within a fixed time window using a weighted exponential decay strategy. (c) Architecture of the BrainMOE module. This module employs a decoupled approach, utilizing subject-specific layers to extract unique information for each subject and subject-shared layers to extract visual features common to all subjects. (d) Inference Pipeline of CrossMind-VL. By performing nearest neighbor search in CLIP space across a video candidate pool, we can retrieve the original video using only brain activity patterns. The copyright of this schematic diagram belongs to the first author of this paper. It should be noted that the natural images included in the schematic are sourced from the open-access dataset CC2017 (https://purr.purdue.edu/publications/2809/1), which is licensed under the CC0 1.0 Universal Public Domain Dedication.

**Figure 2:**
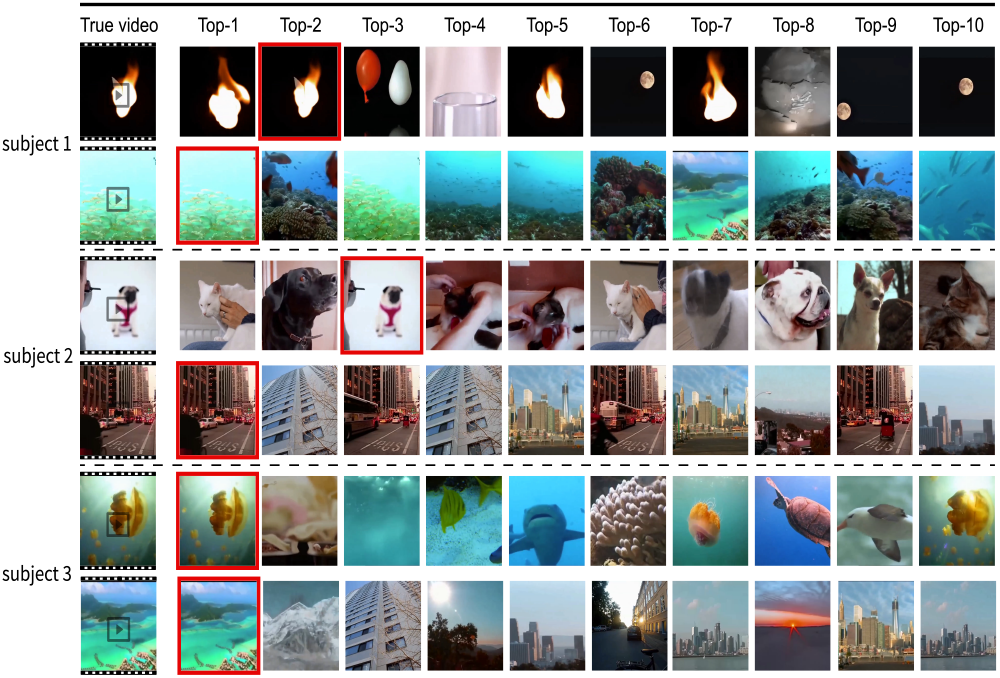
Top-10 retrieval results on the CC2017 dataset. Retrieved videos are highlighted with red borders. It should be noted that the natural images included in the schematic are sourced from the open-access dataset CC2017 (https://purr.purdue.edu/publications/2809/1), which is licensed under the CC0 1.0 Universal Public Domain Dedication.

Additionally, the results indicate that while incorporating data from multiple subjects can enhance model performance to some extent, simple aggregation of multi-subject data fails to achieve optimal performance due to inter-subject variability. This is particularly evident under the ‘Large’ setting, as shown in Table 2, where MS-MLP performed significantly worse than the single-subject model Mind-Animator, and MS-Transformer only matched its performance. In contrast, our proposed BrainMOE significantly outperformed the single-subject model, highlighting the necessity of modeling multi-subject data separately.

### 4.6 New Subject Few-shot Adapting

BrainMOE also demonstrates the ability to transfer pretrained knowledge of shared subject features learned from prior subjects, enabling adaptation to new subjects. This capability is particularly valuable in practical applications, where collecting brain signals for new subjects is both resource-intensive and time-consuming. To simulate scenarios with limited data availability, we evaluated our method using subsets of the full 18-session (4320 samples) training data—specifically, 1, 2, 6, and 10 sessions on the CC2017 dataset for new subject adaptation. Following a leave-one-subject-out paradigm, we selected two subjects for pretraining (source subjects) and one additional subject for adaptation (target subject).

We evaluated different fine-tuning strategies for new subjects, including:

- **Vanilla**: Randomly initializing the BrainMOE model and fine-tuning it solely on the new subject’s data, completely discarding shared features learned from prior subjects.
- **Tune Linear**: Initializing the model with weights pretrained on source subjects, freezing the shared modules, and fine-tuning only the subject-specific linear layer for the new subject.
- **Tune All**: Initializing the model with weights pretrained on source subjects and fine-tuning all parameters for the new subject.

The average results across three experiments are presented in Table 4. As shown in the table, both Tune Linear and Tune All significantly outperform Vanilla in all scenarios, demonstrating that our BrainMOE architecture effectively learns shared features across subjects, enabling adaptation to new subjects with limited data. When only a small amount of data is available for new subjects (1 or 2 sessions), fine-tuning only the linear layer achieves over 90% of the performance of full-parameter fine-tuning while requiring only 23% of the trainable parameters. In some cases, it even surpasses the performance of full-parameter fine-tuning. Furthermore, thanks to the shared features learned by BrainMOE, it achieves over 97% of the performance obtained with the full dataset (24 sessions) using only 10 sessions for new subjects. This advancement holds significant promise for substantially reducing scan times in practical applications, paving the way for more cost-efficient and generalizable brain decoding strategies.

**Table 4:** Few-shot learning accuracy on the CC2017 Dataset. Under the leave-one-subject-out setting, we trained the model on the entire training sets of two other subjects and fine-tuned it on the remaining subject. The training set of the CC2017 dataset comprises 18 sessions, with each session containing 240 samples. We fine-tuned the model on 1, 2, 6, and 10 sessions, respectively, and reported the average accuracy across different subjects.

4.6 New Subject Few-shot Adapting
| Session | Different tune strategy | Parameters | top-10 small | top-100 small | top-10 large | top-100 large |
| --- | --- | --- | --- | --- | --- | --- |
| 1 (5.6%) | Valina | 7.58M | 2.31 | 14.67 | 1.36 | 8.53 |
|  | Ours (tune linear) | 3.46M | 3.17 | 21.83 | 1.89 | 12.42 |
|  | Ours (tune all) | 14.5M | **3.44** | **22.28** | **2.17** | **12.81** |
| 2 (11.1%) | Valina | 7.58M | 2.47 | 17.64 | 1.19 | 8.25 |
|  | Ours (tune linear) | 3.46M | **3.64** | **24.97** | **2.03** | **12.72** |
|  | Ours (tune all) | 14.5M | 3.53 | 24.92 | 1.67 | 12.36 |
| 6 (33.3%) | Valina | 7.58M | 3.97 | 25.08 | 1.78 | 13.06 |
|  | Ours (tune linear) | 3.46M | 4.58 | 29.00 | **3.03** | 15.56 |
|  | Ours (tune all) | 14.5M | **5.08** | **31.31** | 3.00 | **16.25** |
| 10 (55.6%) | Valina | 7.58M | 4.53 | 27.78 | 2.44 | 14.42 |
|  | Ours (tune linear) | 3.46M | 5.14 | 30.00 | **3.25** | 16.06 |
|  | Ours (tune all) | 14.5M | **5.56** | **32.81** | 3.17 | **16.78** |
| Full data | Ours | 14.5M | 5.72 | 32.22 | 3.25 | 17.69 |

### 4.7 Ablation Study and Parameter Sensitivity Analysis

To validate the effectiveness of each component in our model, we conducted comprehensive ablation experiments on the CC2017 dataset. First, to verify the role of the subject-specific layer, we replaced it with a shared linear embedding layer updated using gradients from all subjects (denoted as **w/o Subject Specific Layer**). Second, to assess the contribution of the subject-shared MOE layer, we replaced the MOE module with a standard MLP (denoted as **w/o Subject Shared Layer**). Third, to evaluate the importance of text information extracted by the MLLM for modality alignment, we removed the fMRI-text contrastive loss from the tri-modal contrastive learning objective (denoted as **w/o Text Contrast**). The experimental results, as shown in Table 5, demonstrate that each proposed component is essential for the task. Notably, the MOE module plays a critical role, as its removal leads to a significant performance degradation.

**Table 5:** Ablation study on the different components of our proposed method on the CC2017 dataset. The experimental results were obtained under the setting of the “small test set”.

| Models | Subject1 |  | Subject2 |  | Subject3 |  | Average |  |
| --- | --- | --- | --- | --- | --- | --- | --- | --- |
|  | top-10 | top-100 | top-10 | top-100 | top-10 | top-100 | top-10 | top-100 |
| w/o Subject Specific Layer | 3.21 | 25.13 | 5.02 | 27.65 | 4.25 | 30.18 | 4.16 | 27.65 |
| w/o Subject Shared (MOE) Layer | 3.42 | 22.67 | 4.42 | 26.50 | 3.83 | 26.83 | 3.88 | 25.33 |
| w/o Text Contrast | 4.25 | 26.00 | 5.67 | 31.92 | 4.67 | 32.75 | 4.86 | 30.22 |
| Full Model | **5.00** | **27.92** | **6.17** | **34.83** | **6.00** | **33.92** | **5.72** | **32.22** |

Additionally, we conducted a sensitivity analysis on the hyperparameters used in our model, with the experimental results illustrated in Figure 3. As shown in Figure 3(a), a small number of experts fails to capture shared features across multiple subjects, while an excessive number of experts leads to overfitting, especially when the available data is limited. From Figure 3(b), it is evident that model performance is highly sensitive to the fMRI aggregation time window. A too-small window may miss comprehensive brain responses to the corresponding video stimuli, whereas a too-large window risks incorporating irrelevant brain responses from other stimuli, resulting in a significant performance drop. As depicted in Figure 3(c), the exponential decay rate for fMRI aggregation shows no statistically significant effect on retrieval accuracy. Furthermore, figure 3(d) demonstrates that the balancing term in the loss function significantly impacts retrieval performance. This occurs due to the trade-off between *L*_*Projection*_ and *L*_*Semantic*_ during training: excessive weighting causes the model representations to lose constraints from CLIP text features, while insufficient weighting hinders semantic space alignment through contrastive learning. Both scenarios degrade retrieval accuracy by compromising generalization capability.

**Figure 3:**
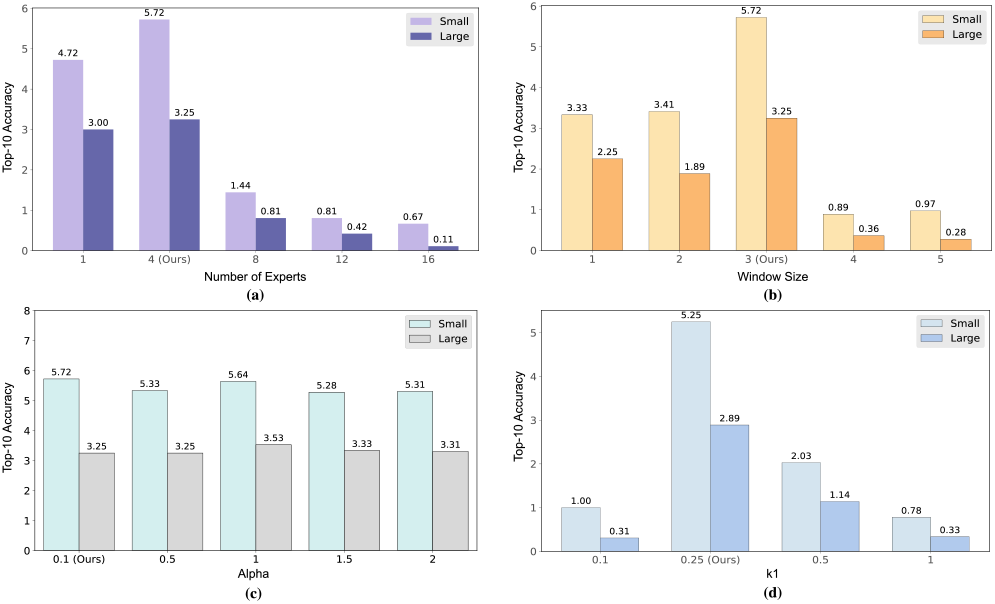
Parameter sensitivity experiments on various hyperparameters were conducted on the CC2017 dataset. Subfigures (a), (b), (c), and (d) illustrate the experimental results for the parameters “number of experts,” “window size,” “alpha,” and “k1,” respectively. All presented results have been averaged across different subjects.

### 4.8 Feature Visualization

To validate the effectiveness of our strategy for separating subject-specific and shared modules, we extracted the output features from both the subject-specific layer and the subject-shared layer of the CC2017 test set and visualized them using t-SNE. As shown in Figure 4, the output features of the subject-specific layer exhibit high separation, demonstrating that this layer effectively captures subject-specific information. In contrast, the output features of the subject-shared layer are clustered together, indicating that this layer successfully extracts common features shared across subjects.

**Figure 4:**
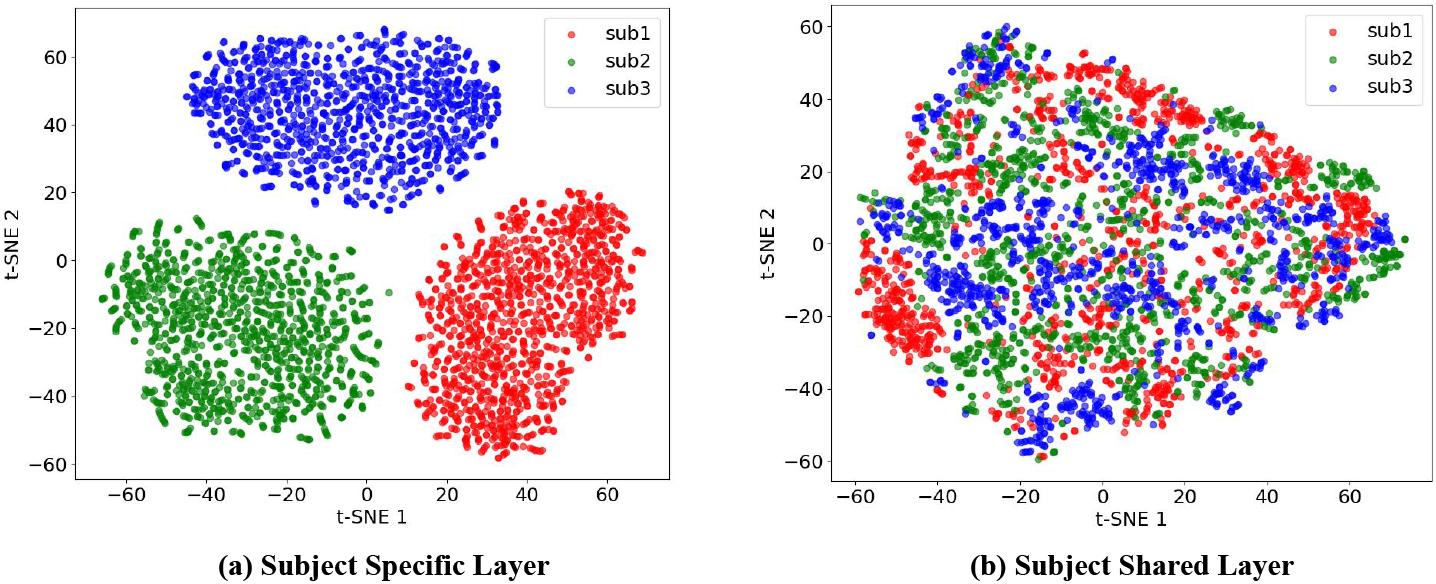
Feature visualizations of different layers in the model. Subfigure (a) displays the features of the subject-specific layer, while subfigure (b) illustrates the features of the subject-shared layer.

## 5 Conclusion

We present a unified framework for cross-subject brain decoding of dynamic visual stimuli that addresses three key challenges: (1) the temporal mismatch between fMRI hemodynamics (0.5 Hz) and video dynamics (30 Hz), (2) the joint modeling of subject-specific and shared neural representations, and (3) the limitations of conventional semantic alignment for dynamic scenes. Our solution introduces: (1) an exponentially weighted multi-frame fusion mechanism for temporal alignment; (2) a hierarchical architecture combining parameterized subject-specific layers with a gated MoE shared representation layer; and (3) a multi-granularity alignment paradigm using Qwen-VL to establish an image-text-brain contrastive learning space.

## References

[1] E. Yargholi and G.-A. Hossein-Zadeh, “Brain decoding-classification of hand written digits from fMRI data employing Bayesian networks,” Frontiers in human neuroscience, vol. 10, p. 351, 2016.

[2] T. Horikawa and Y. Kamitani, “Generic decoding of seen and imagined objects using hierarchical visual features,” Nature communications, vol. 8, no. 1, p. 15037, 2017.

[3] Y. Fujiwara, Y. Miyawaki, and Y. Kamitani, “Modular encoding and decoding models derived from Bayesian canonical correlation analysis,” Neural computation, vol. 25, no. 4, pp. 979–1005, 2013.

[4] K. N. Kay, T. Naselaris, R. J. Prenger, and J. L. Gallant, “Identifying natural images from human brain activity,” Nature, vol. 452, no. 7185, pp. 352–355, 2008.

[5] D. Wildgruber, A. Riecker, I. Hertrich, M. Erb, W. Grodd, T. Ethofer, and H. Ackermann, “Identification of emotional intonation evaluated by fmri,” Neuroimage, vol. 24, no. 4, pp. 1233–1241, 2005.

[6] T. Naselaris, R. J. Prenger, K. N. Kay, M. Oliver, and J. L. Gallant, “Bayesian reconstruction of natural images from human brain activity,” Neuron, vol. 63, no. 6, pp. 902–915, 2009.

[7] R. Beliy, G. Gaziv, A. Hoogi, F. Strappini, T. Golan, and M. Irani, “From voxels to pixels and back: Self-supervision in natural-image reconstruction from fMRI,” Advances in Neural Information Processing Systems, vol. 32, 2019.

[8] N. K. Logothetis, “The neural basis of the blood–oxygen–level–dependent functional magnetic resonance imaging signal,” Philosophical Transactions of the Royal Society of London. Series B: Biological Sciences, vol. 357, no. 1424, pp. 1003–1037, 2002.

[9] S.-G. Kim and S. Ogawa, “Biophysical and physiological origins of blood oxygenation level-dependent fMRI signals,” Journal of Cerebral Blood Flow & Metabolism, vol. 32, no. 7, pp. 1188–1206, 2012.

[10] Y. Lu, C. Du, C. Wang, X. Zhu, L. Jiang, X. Li, and H. He, “Animate your thoughts: Reconstruction of dynamic natural vision from human brain activity,” in The Thirteenth International Conference on Learning Representations.

[11] Z. Chen, J. Qing, and J. H. Zhou, “Cinematic mindscapes: High-quality video reconstruction from brain activity,” Advances in Neural Information Processing Systems, vol. 36, 2024.

[12] A. Vaswani, N. Shazeer, N. Parmar, J. Uszkoreit, L. Jones, A. N. Gomez, L. Kaiser, and I. Polosukhin, “Attention is all you need,” Advances in neural information processing systems, vol. 30, 2017.

[13] J. V. Haxby, J. S. Guntupalli, S. A. Nastase, and M. Feilong, “Hyperalignment: Modeling shared information encoded in idiosyncratic cortical topographies,” elife, vol. 9, p. e56601, 2020.

[14] F. Ozcelik and R. VanRullen, “Natural scene reconstruction from fmri signals using generative latent diffusion,” Scientific Reports, vol. 13, no. 1, p. 15666, 2023.

[15] Z. Chen, J. Qing, T. Xiang, W. L. Yue, and J. H. Zhou, “Seeing beyond the brain: Conditional diffusion model with sparse masked modeling for vision decoding,” in Proceedings of the IEEE/CVF Conference on Computer Vision and Pattern Recognition, 2023, pp. 22 710–22 720.

[16] P. Scotti, A. Banerjee, J. Goode, S. Shabalin, A. Nguyen, A. Dempster, N. Verlinde, E. Yundler, D. Weisberg, K. Norman et al., “Reconstructing the mind’s eye: fmri-to-image with contrastive learning and diffusion priors,” Advances in Neural Information Processing Systems, vol. 36, pp. 24 705–24 728, 2023.

[17] S. Nishimoto, A. T. Vu, T. Naselaris, Y. Benjamini, B. Yu, and J. L. Gallant, “Reconstructing visual experiences from brain activity evoked by natural movies,” Current biology, vol. 21, no. 19, pp. 1641–1646, 2011.

[18] J. Li, D. Li, C. Xiong, and S. Hoi, “Blip: Bootstrapping language-image pre-training for unified vision-language understanding and generation,” in International conference on machine learning. PMLR, 2022, pp. 12 888–12 900.

[19] J. Li, D. Li, S. Savarese, and S. Hoi, “Blip-2: Bootstrapping language-image pre-training with frozen image encoders and large language models,” in International conference on machine learning. PMLR, 2023, pp. 19 730– 19 742.

[20] H. Liu, C. Li, Q. Wu, and Y. J. Lee, “Visual instruction tuning,” Advances in neural information processing systems, vol. 36, pp. 34 892–34 916, 2023.

[21] P. Wang, S. Bai, S. Tan, S. Wang, Z. Fan, J. Bai, K. Chen, X. Liu, J. Wang, W. Ge et al., “Qwen2-vl: Enhancing vision-language model’s perception of the world at any resolution,” arXiv preprint 2409.12191, 2024.

[22] Y. Takagi and S. Nishimoto, “High-resolution image reconstruction with latent diffusion models from human brain activity,” in Proceedings of the IEEE/CVF Conference on Computer Vision and Pattern Recognition, 2023, pp. 14 453–14 463.

[23] Y. Lu, C. Du, Q. Zhou, D. Wang, and H. He, “Minddiffuser: Controlled image reconstruction from human brain activity with semantic and structural diffusion,” in Proceedings of the 31st ACM International Conference on Multimedia, 2023, pp. 5899–5908.

[24] D. Li, C. Du, L. Huang, Z. Chen, and H. He, “Multi-label semantic decoding from human brain activity,” 2018 24th International Conference on Pattern Recognition (ICPR), pp. 3796–3801, 2018. [Online]. Available: https://api.semanticscholar.org/CorpusID:54224318

[25] C. Du, K. Fu, J. Li, and H. He, “Decoding visual neural representations by multimodal learning of brain-visual-linguistic features,” IEEE Transactions on Pattern Analysis and Machine Intelligence, vol. 45, no. 9, pp. 10 760– 10 777, 2023.

[26] J. V. Haxby, M. I. Gobbini, M. L. Furey, A. Ishai, J. L. Schouten, and P. Pietrini, “Distributed and overlapping representations of faces and objects in ventral temporal cortex,” Science, vol. 293, no. 5539, pp. 2425–2430, 2001.

[27] Y. Kamitani and F. Tong, “Decoding the visual and subjective contents of the human brain,” Nature neuroscience, vol. 8, no. 5, pp. 679–685, 2005.

[28] D. Stansbury, T. Naselaris, and J. Gallant, “Natural scene statistics account for the representation of scene categories in human visual cortex,” Neuron, vol. 79, no. 5, pp. 1025–1034, 2013. [Online]. Available: https://www.sciencedirect.com/science/article/pii/S0896627313005503

[29] K. N. Kay, T. Naselaris, R. J. Prenger, and J. L. Gallant, “Identifying natural images from human brain activity,” Nature, vol. 452, pp. 352–355, 2008. [Online]. Available: https://api.semanticscholar.org/CorpusID:54469209

[30] K. Han, H. Wen, J. Shi, K.-H. Lu, Y. Zhang, D. Fu, and Z. Liu, “Variational autoencoder: An unsupervised model for encoding and decoding fmri activity in visual cortex,” NeuroImage, vol. 198, pp. 125–136, 2019. [Online]. Available: https://api.semanticscholar.org/CorpusID:260436433

[31] H. Wen, J. Shi, Y. Zhang, K.-H. Lu, J. Cao, and Z. Liu, “Neural encoding and decoding with deep learning for dynamic natural vision,” Cerebral cortex, vol. 28, no. 12, pp. 4136–4160, 2018.

[32] A. Radford, J. W. Kim, C. Hallacy, A. Ramesh, G. Goh, S. Agarwal, G. Sastry, A. Askell, P. Mishkin, J. Clark et al., “Learning transferable visual models from natural language supervision,” in International conference on machine learning. PMLR, 2021, pp. 8748–8763.

[33] Z. Chen, J. Qing, and J. H. Zhou, “Cinematic mindscapes: High-quality video reconstruction from brain activity,” in Advances in Neural Information Processing Systems, A. Oh, T. Naumann, A. Globerson, K. Saenko, M. Hardt, and S. Levine, Eds., vol. 36. Curran Associates, Inc., 2023, pp. 24 841–24 858.

[34] J. V. Haxby, J. S. Guntupalli, S. A. Nastase, and M. Feilong, “Hyperalignment: Modeling shared information encoded in idiosyncratic cortical topographies,” eLife, vol. 9, p. e56601, jun 2020. [Online]. Available: 10.7554/eLife.56601

[35] P.-H. C. Chen, J. Chen, Y. Yeshurun, U. Hasson, J. Haxby, and P. J. Ramadge, “A reduced-dimension fmri shared response model,” in Advances in Neural Information Processing Systems, C. Cortes, N. Lawrence, D. Lee, M. Sugiyama, and R. Garnett, Eds., vol. 28. Curran Associates, Inc., 2015. [Online].

[36] W. Li, M. Liu, F. Chen, and D. Zhang, “Graph-based decoding model for functional alignment of unaligned fmri data,” Proceedings of the AAAI Conference on Artificial Intelligence, vol. 34, pp. 2653–2660, 04 2020.

[37] N. Shazeer, A. Mirhoseini, K. Maziarz, A. Davis, Q. Le, G. Hinton, and J. Dean, “Outrageously large neural networks: The sparsely-gated mixtureof-experts layer,” arXiv preprint 1701.06538, 2017.

[38] D. S. Marcus, J. Harwell, T. Olsen, M. Hodge, M. F. Glasser, F. Prior, M. Jenkinson, T. Laumann, S. W. Curtiss, and D. C. Van Essen, “Informatics and data mining tools and strategies for the human connectome project,” Frontiers in neuroinformatics, vol. 5, p. 4, 2011.

[39] K. Han, H. Wen, J. Shi, K.-H. Lu, Y. Zhang, D. Fu, and Z. Liu, “Variational Autoencoder: An unsupervised model for encoding and decoding fMRI activity in visual cortex,” NeuroImage, vol. 198, pp. 125–136, 2019.

[40] M. F. Glasser, T. S. Coalson, E. C. Robinson, C. D. Hacker, J. Harwell, E. Yacoub, K. Ugurbil, J. Andersson, C. F. Beckmann, M. Jenkinson et al., “A multi-modal parcellation of human cerebral cortex,” Nature, vol. 536, no. 7615, pp. 171–178, 2016.

[41] C. Wang, H. Yan, W. Huang, J. Li, Y. Wang, Y.-S. Fan, W. Sheng, T. Liu, R. Li, and H. Chen, “Reconstructing rapid natural vision with fMRI-conditional video generative adversarial network,” Cerebral Cortex, vol. 32, no. 20, pp. 4502–4511, 2022.

[42] G. Kupershmidt, R. Beliy, G. Gaziv, and M. Irani, “A penny for your (visual) thoughts: Self-supervised reconstruction of natural movies from brain activity,” arXiv preprint 2206.03544, 2022.

